# Mapping the human sperm proteome across compartments, donors and individual cells

**DOI:** 10.64898/2026.09.22.753411

**Authors:** Leander van der Hoeven, Luisa Schmidt, Søren Ziebe, Morten Rønn Petersen, Kristian Almstrup, Tanveer S. Batth, Anders Rehfeld, Jesper V. Olsen

## Abstract

The human spermatozoon is among the most specialised cells in the body, yet where its proteins reside, where they come from and how they vary between and within men remain largely unresolved. Here we present the deepest proteomic atlas of human spermatozoa to date, identifying 10,828 proteins across six donors. Abundances correlate well between individuals and are anchored by a conserved core of mitochondrial energy-metabolism proteins, while a smaller fraction varies between donations from the same man. By mechanically separating heads from tails, we assigned thousands of previously unlocalised proteins to a compartment, and by comparing seminal fluid from vasectomised and non-vasectomised men we distinguished sperm-and epididymis-derived proteins from accessory-gland secretions. Using a single-cell analysis workflow we further show that the proteomes of individual spermatozoa can be measured, recovering up to ∼1,000 protein groups per cell. In a cohort with total fertilisation failure, we find recurrent mitochondrial rather than sperm–egg fusion dysregulation. Together or data provides novel insights into the dynamics of the sperm proteome and male infertility

## Introduction

Human sperm cells are among the most specialised cell types in the body, both morphologically and functionally, uniquely serving the purpose of fertilisation (1). They are also of growing clinical concern: approximately one in six couples experience difficulty conceiving, and meta-analyses report a substantial decline in average sperm counts over the past five decades (2, 3, 4). Despite this, there is no targeted treatment for male infertility besides for cases with obstruction or infection, and diagnosis still rests on conventional semen analysis, which describes cells rather than explaining their function/dysfunction (5, 6). Molecular characterisation is complicated by the nature of the cell itself. Sperm cells were long thought to lack active protein production and to be transcriptionally silent (7). More recent studies show that, although transcriptionally restricted, they are not completely silent (8, 9), as mitochondrial translation can occur (10, 11, 12).

Infertility research still lacks a fully comprehensive proteomics-based resource for sperm (5, 13), despite a few published studies (1, 5, 13, 14, 15). Early studies relied on two-dimensional gel electrophoresis with limited mass-spectrometric identification, reporting only 100–260 proteins (16–18). The advent of high-sensitivity liquid chromatography tandem mass spectrometry (LC-MS/MS)-based shotgun proteomics raised this to approximately 1,760 proteins in the first large-scale sperm proteome (19), challenging the assumption that spermatozoa have a relatively simple protein composition (20). Further advancements in mass-spectrometry sensitivity have since pushed this further: recent experimental studies report close to 10,000 proteins (13, 15), while computationally combining multiple studies suggests up to 9,296 proteins may be present (14). The most recent of these, by Greither et al. and Kong et al. (21, 15), have brought coverage of the sperm proteome close to that achieved in other human tissues. Despite these advances (1, 14), technical limitations mean the sperm proteome likely remains incomplete (5). Depth alone, however, says little about where these proteins reside within the cell, which of them are contributed by the tissues producing the surrounding fluid, or how consistent they are between and within men, which determine whether a protein catalogue can be interpreted biologically. Due to the characteristic of sperm cells itself, analysis has also been more challenging than for somatic cells. Sperm cells are morphologically unusual with densely packed chromatin (22), and seminal plasma proteins remain among the most abundant proteins detected even after the plasma is removed prior to LC-MS/MS (5). Seminal fluid is similarly understudied relative to its importance (23, 24). Its proteins nourish and protect sperm cells and interact with the female reproductive tract to influence fertilisation outcomes (23, 25, 26), and alterations in the seminal plasma proteome, including increased annexin A2 (ANXA2) and reduced semenogelin-2 (SEMG2), as well as changes in oxidative-stress-associated proteins such as DJ-1, lactotransferrin and peroxiredoxins, have been associated with infertility (26–28). Whether these alterations are causal remains unknown (23, 24), in part because the tissue of origin of most seminal plasma proteins has not been systematically resolved. Seminal plasma receives contributions from the testis, epididymis, prostate, seminal vesicles and other glands, together with proteins shed from the urogenital epithelium (5, 23). Resolving these contributions would indicate whether an abnormal seminal proteome reflects the sperm cells themselves or the tissues supplying the fluid around them, a distinction with direct diagnostic consequences and potential therapeutic options.

Another limitation in regard to sperm analysis is that current approaches do not resolve where in the cell the identified proteins reside, a dimension that is crucial for interpreting sperm function and thereby can potentially be important for infertility diagnostics. Sperm cells are highly compartmentalised: the head contains the nucleus and acrosome and mediates binding to and penetration of the zona pellucida, while the flagellum is joined to the head by a neck region and divided into midpiece, principal piece and end piece, with the mitochondria that power motility housed in the midpiece (29, 30, 31). For most sperm proteins, the compartment in which they reside is unknown (5, 29). Because function in this cell type is so tightly coupled to location, assigning proteins to compartments is a prerequisite for interpreting them (29, 32).

Here we combine deep bulk proteomics, mechanical head-tail separation, single-cell analysis and seminal fluid profiling to describe the human sperm proteome and how it is organised. We identify 10,828 proteins across six donors and assess how stable these abundances are between men and across repeated donations from the same man. Separating heads from tails allows thousands of previously unlocalised proteins to be assigned to a compartment, while an automated single-cell workflow establishes that proteomic measurement of individual spermatozoa is feasible despite their 2 to 5 pg protein content. Comparing seminal fluid from vasectomised and non-vasectomised men distinguishes testis-and epididymis-derived proteins from accessory gland secretions and traces their tissue of origin. Finally, in men with total fertilisation failure, we find recurrent mitochondrial dysregulation rather than the sperm-egg fusion defects anticipated.

## Material and Methods

### Sample preparation of swim-up sperm samples

Semen samples were obtained from six self-reported healthy donors, who were instructed to abstain from ejaculation for at least 48 h prior to sample delivery, by masturbation into sterile wide-mouthed plastic containers at the Department of Growth and Reproduction, Rigshospitalet, and allowed to liquefy at 37°C for 15 to 30 minutes prior to processing. Donors received a reimbursement of 500DKK per sample for their inconvenience. Spermatozoa were isolated from liquefied ejaculates by swim-up into human tubal fluid medium (HTF) composed of 97.8 mM NaCl, 4.69 mM KCl, 0.2 mM MgSO₄, 0.37 mM KH₂PO₄, 2.04 mM CaCl₂, 0.33 mM sodium pyruvate, 21.4 mM sodium lactate, 2.78 mM glucose, 21 mM HEPES, and 4 mM NaHCO₃, with the pH adjusted to 7.4 using NaOH. Following a 1-hour incubation at 37 °C, the swim-up fraction was collected into a separate tube and centrifuged at 700g for 10mins, whereafter the sperm concentration was measured by image cytometry (33). Following this, the sample was washed with HTF and centrifuged at 700g for 10 mins four times before the sperm pellet was flash frozen using liquid nitrogen and stored at-80°C until analysis.

Thawed sperm pellets were washed twice with ice-cold HTF buffer by centrifuging at 600 xg for 5 min. For cell lysis, the sperm cells were lysed with 1 mL of boiling SDS lysis buffer (4% SDS; 100 mM Tris pH 8.5, 5 mM TCEP, and 10 mM CAA), added directly to the cell pellet, followed by incubation at 95 °C for 10 min with mixing (1000 rpm). Lysates were sonicated with a tip probe (1 minute, 1 second on, 2 seconds off, 60% amplitude, Fisherbrand™ probes Model 120 Sonic Dismembrator). Protein concentration was calculated using the BCA assay and tryptophan assay. Following this 50µg of protein Lysate was digested per sample. Digestion was done following the Protein Aggregation Capture (PAC) protocol as previously described by Batth et al. (34). Briefly, proteins were aggregated on magnetic microbeads (MagResyn Hydroxyl beads) by adding acetonitrile (ACN) to a final concentration of 70%. Beads were washed once with 100% ACN and once with 70% ethanol. 50 mM HEPES digestion buffer at pH 8.5 was added to all samples prior to the addition of proteases. Unless stated otherwise, a typical protease-to-protein ratio of 1:100 was employed for digestion. The digestion reaction was quenched by adding trifluoroacetic acid (TFA) to a final concentration of 1%. Thereafter, an amount equivalent to 700 ng of the digestion product was loaded on Evotips for subsequent MS analysis.

### Head-Tail separation

Sperm pellets from the same donors as above were combined and washed as previously described. After the washing, the combined cell pellet of the six donors was resuspended into 200 uL of HTF buffer. After resuspension, cells were sonicated using the 120 Sonic Dismembrator and a tip probe for 25 seconds, 1 second on/2 seconds off, and at 60% amplitude. The cell suspension was centrifuged at 750 xg for 1 min. Subsequently, the pellet (head fraction) was lysed using SDS lysis buffer for 10 min at 95 °C. Further, the supernatant was centrifuged at 5000 xg for 3 min, and the pellet (tail fraction) was handled as described above. The head and tail protein lysates were subsequently sonicated again as described in the sample preparation section and digested using PAC. Peptides from the head and tail samples were combined in various fractions (0:100, 20:80, 40:60, 60:40, 80:20 and 100:0), with all of the combined samples having an input of around 700 ng and loaded on Evotips for subsequent MS analysis.

### Seminal Fluid

Seminal fluid samples were collected at the Department of Growth and Reproduction, Rigshospitalet. Semen samples from vasectomised and non-vasectomised men were centrifuged at 3000 xg for 30 min, and the supernatant (seminal fluid) was kept for further analysis. Only seminal fluid samples from vasectomised men with a confirmed absence of sperm cells were included. Absence of sperm was verified by cytological examination of the pellet by microscopy. Following this, the pellet was flash frozen using liquid nitrogen and stored at-80°C. Seminal fluid samples were centrifuged at 700 xg for 5 min prior to handling steps. SDS lysis buffer was added in equal volume and incubated for 10 min at 95 °C with mixing (1000 rpm). Samples were sonicated and prepared for MS as described above.

### Total fertilisation failure Cohort

Men were recruited by recall in an earlier study (35) based on a total failure of fertilisation in IVF attempts. Briefly, in the selection of the TFF group, no zygotes developed after IVF despite oocytes and semen parameters appearing normal, and the inclusion criteria were as follows: (i) ≥ 4 oocytes retrieved for IVF treatment; (ii) sperm concentration ≥ 15 × 10^6^ cells/ml; and (iii) successful fertilisation in a subsequent ICSI treatment. The exclusion criteria were as follows: (i) couples with known female or male causes of infertility affecting oocyte or semen quality, and (ii) couples with donor cycles. In the selection of the control group, the inclusion criteria were as follows: (i) ≥ 4 oocytes retrieved for IVF treatment; (ii) sperm concentration ≥ 15 × 10^6^ cells/ml; and (iii) a fertilisation rate of 60% or more. They were matched according to the female age in the TFF group (±2 years) and were selected among patients receiving treatment two months prior to or after the couples experiencing TFF. Controls were also selected to match the oocyte, and sperm counts of the TFF group, but this was not always possible. From the original TFF cohort, 5 out of 17 TFF men and 8 out of 16 controls were included in this study, based on sample availability.

Semen parameters were recorded as part of the original study (35). In brief, the men from the recruited couples were instructed to abstain from ejaculation for at least 48 h prior to sample delivery. Samples were produced by masturbation, collected into sterile wide-mouthed plastic containers, and allowed to liquefy before being mixed thoroughly for analysis. Semen volume was determined by weight, and the sperm concentration measured by image cytometry (33). Motility was scored by applying 10 μl of semen to a pre-warmed (37°C) glass slide and immediately examining it on a heated (37°C) microscope stage, with spermatozoa classified as progressively motile or non-progressive/immotile. For morphology, semen smears were Papanicolaou stained and independently evaluated by two experienced technicians. Total progressive motile sperm count (TPMSC) was calculated as the product of progressive motility (%) and sperm concentration. Because of the limited number of spermatozoa per ejaculate, not all parameters could be assessed in every sample. Raw semen aliquots of 200 μl were stored at −80°C and eventually provided for this study.

### Single-sperm analysis

Sperm cells from swim-up samples were pelleted by centrifugation in a swing-out centrifuge at 800 × g for 3 min and washed three times with degassed PBS. Cells were diluted to approximately 200 cells/µL in PBS and kept on ice until single-cell sorting. Single-cell isolation and digestion were performed using the cellenONE (Cellenion). First, 300 nL of ice-cold master mix was dispensed into each well of LoBind 96-well plates (Eppendorf) using a PDC M Piezo Dispensing Capillary (PDC; Cellenion) with an average droplet volume of approximately 325 pL. The master mix consisted of 30 mM triethylammonium bicarbonate (TEAB, pH 8.5), 0.02% n-dodecyl-β-D-maltoside (DDM), 10 ng/µL Trypsin Gold (Promega, Mass Spectrometry Grade), 5 ng/µL Lys-C (Wako), and 250 µM dithiothreitol (DTT, Proteomics Grade). The master mix was kept on ice, mixed thoroughly, and centrifuged at 14,000 × g for 2 min immediately before aspiration and dispensing. After dispensing the master mix, the PDC was washed twice with 180 µL of 8 M urea, followed by two flushes to remove residual master mix. Next, the cellenONE was used to characterize the properties of the cell population, and individual sperm cells (n = 76) with diameters of 5–8 µm and a maximum elongation factor of 4× were isolated into wells containing the master mix. In addition, for each donor, pools of 0, 5, 10, or 20 sperm cells (n = 5 wells per condition) were dispensed into individual wells. Following cell dispensing, the PDC was again washed twice with 180 µL of 8 M urea, followed by two flushes to remove residual cell debris. Both master mix dispensing and cell isolation were performed at 8 °C and 65% relative humidity. Digestion was subsequently carried out on the cellenONE at 37 °C and 85% relative humidity for 2.5 h. During digestion, the PDC continuously dispensed 300 pL of distilled water into each well to compensate for evaporation and prevent sample drying. The digestion reaction, with a final volume of approximately 2 µL, was quenched and stabilized by the addition of 1 µL of 5% formic acid, 2% acetonitrile. Plates were briefly centrifuged at 250 xg to collect all liquid at the bottom of the wells and were stored at 7 °C for up to one week before subsequent MS analysis.

### LC-MS/MS analysis of single sperm cells

Single-sperm samples were analysed using an Orbitrap Astral Zoom mass spectrometer equipped with a Nanospray Flex™ Ion Source and a Thermo Scientific™ FAIMS Pro Duo Interface (Thermo Scientific). The spray voltage was set to 2 kV, the capillary temperature to 275 °C, and the RF level to 50%. Full precursor (MS1) scans were acquired using the Orbitrap™ mass analyzer, whereas tandem fragment (MS2/Data-Independent Acquisition (DIA)) scans were acquired in parallel using the Astral™ mass analyzer. Orbitrap lock-mass correction was performed at the start of each run using EASY-IC. The FAIMS compensation voltage (CV) was set to −46 V for all scans. For data acquisition, pre-accumulation and low-input mode were enabled. MS1 scans were acquired over an m/z range of 400–800 with a resolution of 240,000, a normalised Automatic Gain Control (AGC) target of 500 (5,000,000 charges), and a maximum injection time of 25 ms. DIA scans were acquired over the same 400–800 m/z precursor range using 10 m/z isolation windows with a maximum injection time of 20 ms. Fragmentation was performed by higher-energy collisional dissociation (HCD) using a normalised collision energy (NCE) of 25. Fragment ions were detected over an m/z range of 150–1,500 with a normalised AGC target of 200 (20,000 charges). Loop control was set to “All”, resulting in a combined MS1/DIA cycle time of approximately 1s.

### LC-MS/MS analysis of bulk sperm samples

The peptide samples were eluted from Evotips using an Evosep One LC system (Evosep Biosystems) and separated using an Evosep 8 cm (EV1109, Evosep) performance column connected to a steel emitter (EV1086, Evosep) and heated to 40 °C over a 21 min (60 SPD) gradient or 42 min (30SPD) gradient. The eluted peptides were electrosprayed into an Orbitrap Astral mass spectrometer (Thermo Fisher Scientific) using 1800 V spray voltage, funnel radio frequency level at 40, and a heated capillary temperature set to 275 °C in positive mode. Full scan precursor spectra (380–980 Da) were recorded in profile mode using a resolution of 240,000 at 200 m/z, a normalized automatic gain control (AGC) target of 500%, and a maximum injection time of 3 ms. The fragment spectra were acquired in narrow-window data-independent acquisition (nDIA) mode as previously described (36), with a precursor mass range of 380 to 980 m/z with 2 Thomson isolation windows. Isolated precursors were fragmented in the HCD cell using 25% normalized collision energy, a normalized AGC target of 500%, and a maximum injection time of 2.5 ms.

## Data analysis

The MS RAW files were analysed using either SpectronautV21 or DIANN v2.5.6 (academia) searching against a FASTA file containing the human proteome (SwissProt, 20,421 sequences, downloaded on May 6th, 2025) using the default search settings. In brief, carbamidomethyl was set as a fixed modification for cysteine and N-terminal acetylation and oxidation of methionine as variable modifications. Trypsin was specified as the proteolytic enzyme with a maximum of one missed cleavage allowed. The mass tolerance was set to automatic inference at both MS1 and MS2 levels with a precursor false discovery rate (FDR) level at 1%. Quantification was performed using the Quant UMS (high precision) setting. For single cell analysis we excluded injection with no protein identification, removed proteins identifications that were not enriched over the blanks and required quantification in at least 75% of the samples per group. The data was analysed in R (v4.5.1) with RStudio (v2025.05.1) using the following packages: tidyverse (v2.0.0), dplyr (v1.1.4), tidyr (v1.3.1), tibble (v3.3.0), stringr (v1.5.2), purrr (v1.1.0), readr (v2.1.5), forcats (v1.0.1), data.table (v1.17.8), arrow (v21.0.0.1), readxl (v1.4.5), writexl (v1.5.4), limma (v3.64.3), clusterProfiler (v4.16.0), enrichplot (v1.28.4), DOSE (v4.2.0), fgsea (v1.34.2), GOSemSim (v2.34.0), msigdbr (v25.1.1), org.Hs.eg.db (v3.21.0), GO.db (v3.21.0), AnnotationDbi (v1.70.0), biomaRt (v2.64.0), irlba (v2.3.5.1), matrixStats (v1.5.0), mgcv (v1.9-3), MASS (v7.3-65), ggplot2 (v4.0.0), ggrepel (v0.9.6), patchwork (v1.3.1), ggfortify (v0.4.19), ggpubr (v0.6.2), ggVennDiagram (v1.5.4), scales (v1.4.0), pheatmap (v1.0.13), RColorBrewer (v1.1-3), UpSetR (v1.4.0), corrplot (v0.95), factoextra (v1.0.7), gridExtra (v2.3), igraph (v2.1.4), httr (v1.4.7), jsonlite (v2.0.0), future.apply (v1.20.0), doParallel (v1.0.17), BiocManager (v1.30.26) and conflicted (v1.2.0).

## Results

### A comprehensive, validated atlas of the human sperm proteome

To generate a comprehensive reference atlas of the human sperm proteome, which can also serve as a deep spectral library, we isolated sperm cells from six donors and analysed them using three approaches: pooled single-shot nDIA, per-donor single-shot analysis combined post-acquisition, and off-line high-pH reversed-phase fractionation (46 fractions). We employed 3 methods to determine first, all the proteins that can be detected through the high-pH reversed-phase fractionation. The other two methods were employed to determine how many proteins we could detect from single-shot injections. All were measured on an Orbitrap Astral coupled to an Evosep Eno LC (nDIA, 30 SPD). The pooled samples yielded approximately 9,300 unique proteins, per-donor analysis over 10,000, and fractionation almost 11,000 proteins (Figure 1A), with peptide-per-protein counts indicating excellent coverage (Supplementary Figure 1A). In total, we identified approximately 185,000 precursors (a peptide in one specific charge and modification state) corresponding to around 150,000 peptides (Supplementary Figure 1B). We compared our dataset to two previously published sperm proteome studies, by Greither et al. and Kong et al., the latter the most recent (21, 15), identifying more protein groups than either. To confirm this was not simply divergence from the literature, we calculated the Pearson correlation of relative protein intensities against the Kong dataset (8,084 shared proteins), obtaining r = 0.855 (Figure 1B, Supplementary Figure 1C), a high value given the different donors and acquisition methods used, in contrast, the correlation to Greither et al. was substantially lower (r = 0.589). Rank plots of median protein intensity tracking the top 10 most abundant proteins of our study across the published datasets showed that most proteins retained a comparable relative rank, with some shifting toward a more median rank in the other studies (Figure 1C). Together, these comparisons indicate that our dataset reproduces the results of the previously published sperm proteome, and that the additional protein groups therefore represent increased analytical depth rather than a systematically divergent measurement.

**Figure 1.**
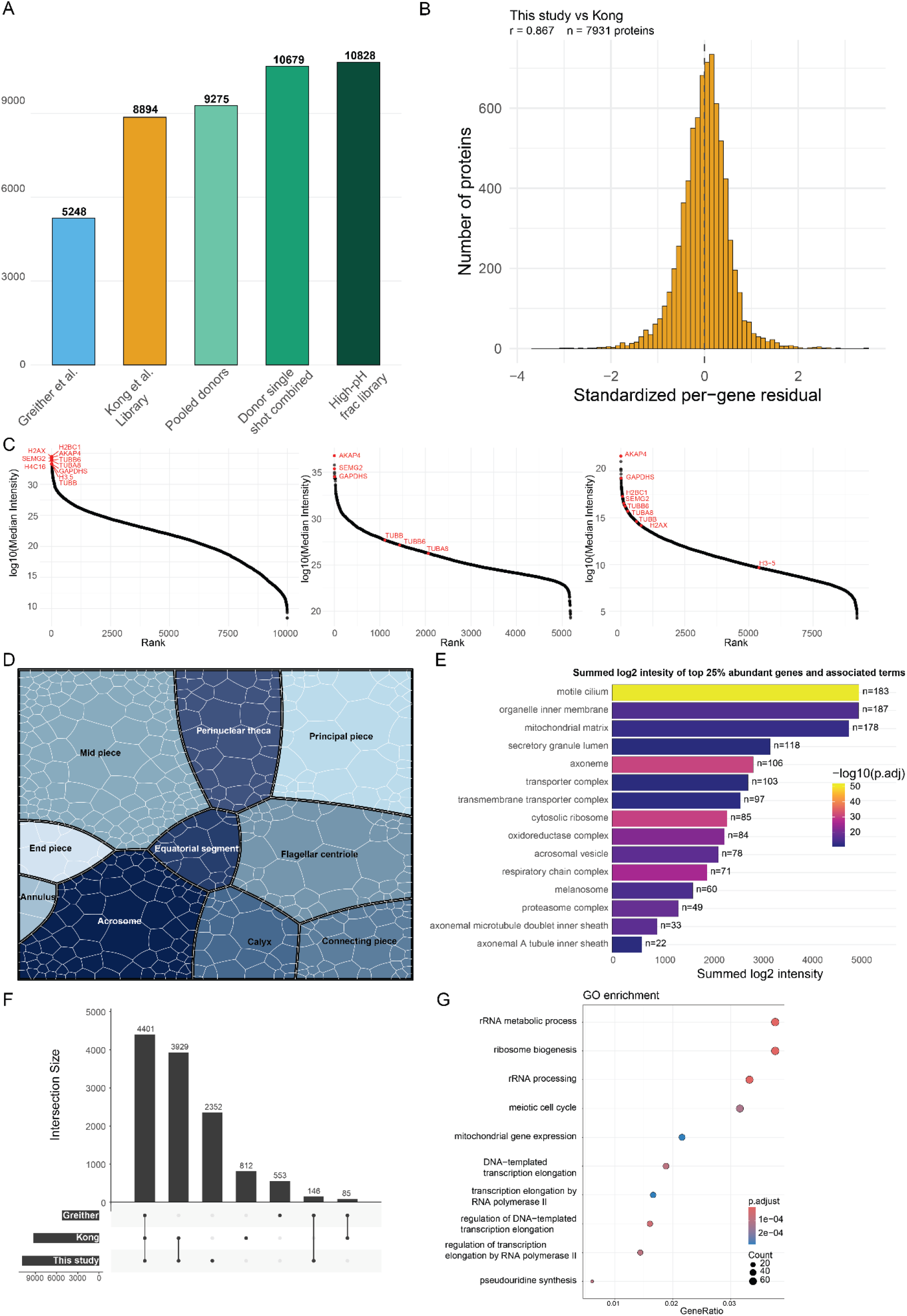
Depth and validation of the human sperm proteome. (A) Number of unique protein groups identified per approach: previously published datasets (Greither et al., Kong et al.) shown for reference, alongside pooled single-shot analysis, per-donor single-shot analysis combined post-acquisition, and high-pH reversed-phase fractionation (46 fractions) of this study. (B) Distribution of standardized per-protein residuals between this study and Kong et al., restricted to the 7,545 proteins detected in both datasets. (C) Rank plots of median protein intensity for this study and previously published datasets. The top 10 most abundant proteins of this study are highlighted (red) and tracked across the other datasets. (D) Voronoi treemap of sperm proteins assigned to subcellular compartments using Human Protein Atlas localisation annotations. Each cell represents one protein, sized by its intensity (E) Significantly enriched Gene Ontology (GO) cellular-component terms among the 25% most abundant proteins, ranked by summed log2 protein intensity. Bar colour denotes the adjusted enrichment *P* value, and labels indicate the number of proteins annotated to each term. (F) UpSet plot of protein overlap between this study, Kong et al. and Greither et al., showing 2,655 proteins unique to this study. (G) GO term enrichment of the proteins newly identified in this study relative to Kong et al., using all the other proteins detected as background

We next examined how protein abundances are distributed across sperm compartments. Using the Human Protein Atlas (HPA) subcellular annotations (37), we created a Voronoi treemap in which each protein is drawn as a cell whose area is proportional to its intensity (Figure 1D). We also performed gene ontology (GO) enrichment on the top 25% most abundant proteins (Figure 1E). The midpiece proteins accounted for the largest share of total abundance, followed by the acrosome, flagellar centriole, and principal piece. The prominence of the acrosome and principal piece likely reflects that these are the most-studied compartments. The flagellar centriole, despite its small area, illustrates how a compartment can disproportionately represent protein abundance and function (Figure 1D). This emphasis on the flagellum is reinforced in Figure 1E, where the most abundant proteins are associated with the cilium and mitochondria. We identified 2,655 proteins unique to our dataset relative to Kong et al. (Figure 1F). GO term enrichment analysis of these newly identified proteins showed the strongest overrepresentation for chromosome segregation, DNA replication and the meiotic cell cycle, together with DNA repair and rRNA processing (Figure 1G).

To further benchmark completeness, we compared our dataset to the theoretical sperm proteome that Pini et al. derived by computationally combining detected proteins from several sperm proteomics studies (14). Our dataset recovered 85% of these proteins, with only 900 unique to Pini et al. and over 4,800 unique to ours (Supplementary Figure 1D), covering most of the previously reported sperm proteome while adding substantially to it. Because swim-up isolation reduces but does not fully eliminate contamination by other cell types (38), we next assessed whether proteins from organelles not expected in the mature spermatozoon were overrepresented (Supplementary Figure 1E). ER and Golgi proteins were detected at relatively high intensity, and the abundance ranks of the most highly expressed HeLa proteins were distributed across the rank range, although a subset appeared at high rank (Supplementary Figure 1F). Both patterns closely resemble those of the published datasets, which were generated with different preparations and instruments, arguing against a contaminant specific to our workflow. They do not, however, distinguish proteins genuinely retained in sperm from a low-level contribution common to all sperm preparations.

### Inter-donor variability and temporal stability of the sperm proteome

We next assessed inter-donor variability. The number of proteins identified was remarkably consistent across donors, ranging from 9,844 to 10,005 (Figure 2A), though PCA revealed that some donors differed substantially while others clustered closely (Supplementary Figure 2A). Clustering the most abundant proteins by log2 intensity and identifying the top 5 enriched GO terms per cluster (Figure 2B) showed that the most abundant proteins were again cilium, macro-autophagy, and vesicle proteins, consistent with the relatively silent sperm cell relying on autophagy and vesicles rather than active synthesis machinery. The lowest-intensity proteins were associated with ribosomes and transcription, consistent with the autophagic elimination of ribosomes during spermiogenesis (39), which leaves only residual amounts of the biosynthetic machinery in the mature cell. We then calculated the inter-donor coefficient of variation (CV), classifying proteins detected in all donors with CV below 2.5% as highly conserved, those above 2.5% as moderately conserved, and those not detected in all donors as not universally present, yielding 385, 9,680 and 129 proteins, respectively (Figure 2C).

**Figure 2.**
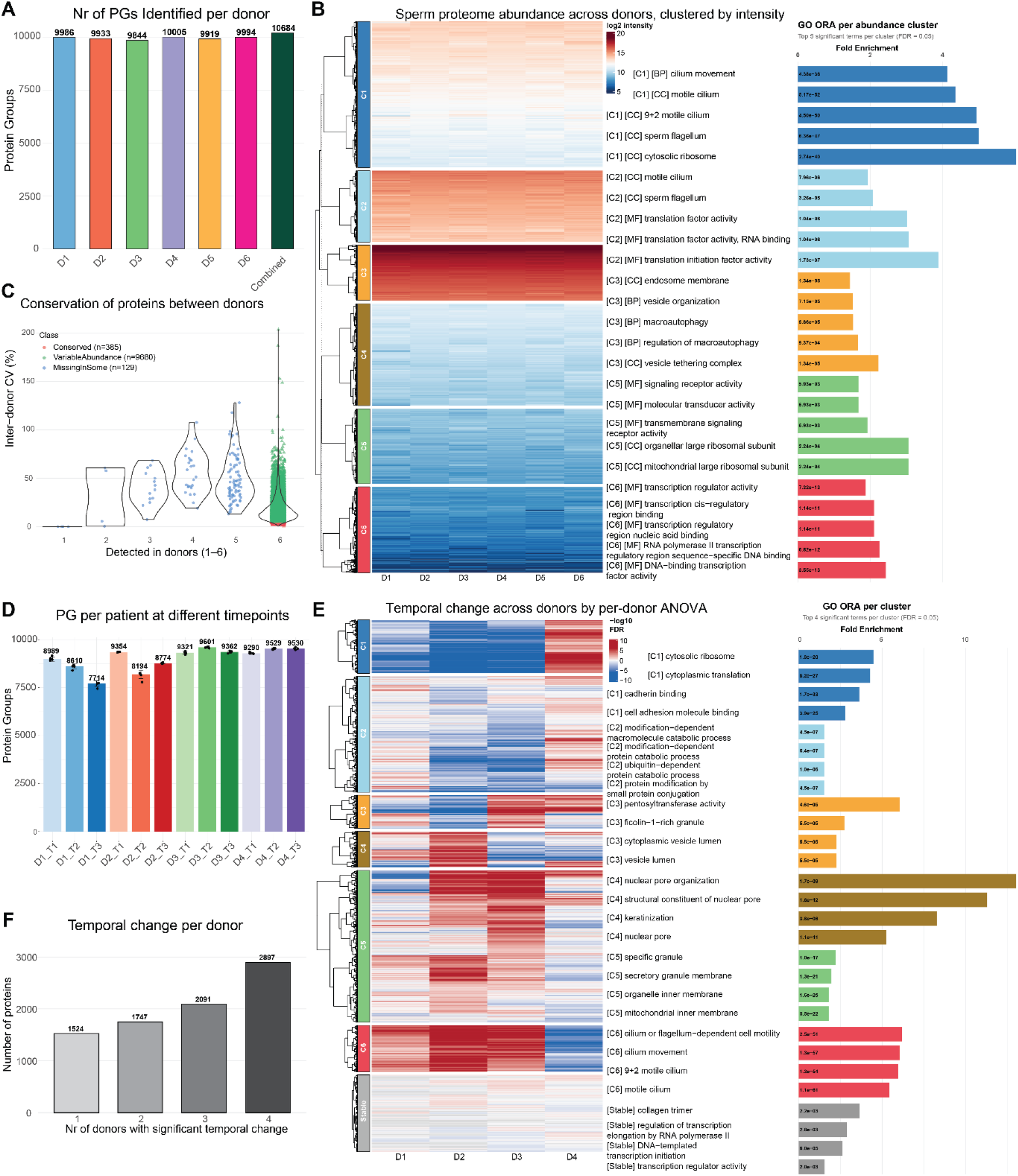
Inter-donor variability and temporal stability of the sperm proteome. (A) Number of protein groups identified per donor across six healthy individuals (range 9,844–10,005) and in the combined set (10,684). (B) Heatmap of log2 protein intensities across donors, hierarchically clustered. The top 5 enriched GO terms per abundance cluster are shown on the right (fold enrichment; FDR < 0.05). (C) Classification of proteins by inter-donor reproducibility: conserved (CV < 2.5%, detected in all donors; n = 385), variable abundance (CV > 2.5%, detected in all donors; n = 9,680), and missing in some donors (n = 129). (D) Number of protein groups per donor at different donation time points (T1–T3) for four donors. (E) Heatmap of per-donor temporal change across time points based on ANOVA, hierarchically clustered, with the top enriched GO terms per cluster shown on the right (fold enrichment; FDR < 0.05). (F) Number of proteins showing significant temporal change, as a function of the number of donors in which the change was significant (1,524–2,897).

To characterise these conserved proteins functionally, we ran STRING association network analysis on the top 100. Nearly all formed a single, highly interconnected cluster (Supplementary Figure 2B), enriched for cellular respiration, ribonucleotide biosynthesis, ATP metabolism and oxidative phosphorylation (Supplementary Figure 2C). This points to mitochondria as the primary source of the most conserved proteins, as expected given the central role of oxidative phosphorylation in sperm motility. The least conserved proteins, by contrast, did not form a coherent network (Supplementary Figure 2D) and were enriched for vesicle and extracellular-space terms (Supplementary Figure 2E). Interestingly, these terms are associated with post-testicular maturation and are epididymis-derived (40, 41), suggesting that the acquisition of such proteins via epididymosomes during epididymal transit may account for their greater variability across donors. We next assessed temporal stability across donation time points in four of the six donors with repeated samples available. Given that the sperm RNA landscape is known to change rapidly with diet and abstinence (42, 43), we examined whether the proteome showed comparable variability. Two donors’ sperm proteomes remained relatively stable, while the other two showed protein-count drops of approximately 14% (8,989 to 7,714) and 12% (9,354 to 8,194) between time points (Figure 2D). These drops reflect genuine turnover (Supplementary Figure 3C). PCA showed minimal separation between time points for the stable donors and pronounced separation for the variable donors (Supplementary Figure 3A), with correspondingly tighter or broader CV distributions (Supplementary Figure 3B).A one-way analysis of variance (ANOVA) across donor×time-point conditions (Benjamini–Hochberg FDR < 0.05) identified proteins varying most across these conditions, which clustered into groups distinguished by the time points at which they were elevated or reduced (Figure 2E); the number of proteins showing significant temporal change ranged from 1,524 to 2,897 depending on the number of donors involved (Figure 2F). Representative trajectories illustrate the range of temporal behaviours: shared increase (e.g. RNASET2), shared decrease (e.g. ALDH1L1), donor-specific change (LTF), and stability (ABCA3) (Supplementary Figure 3D). The 3,408 proteins that did not differ significantly across time points were enriched for cilium and flagellar structural components and ribosomal translation machinery (Supplementary Figure 3E).

### Subcellular resolution of the sperm proteome into head and tail compartments

Mature sperm cells comprise three main functional areas: the head, the midpiece (part of the neck/tail complex), and the tail. To spatially resolve the sperm proteome, we developed an approach to assign proteins to the head or tail compartment, respectively (Figure 3A). Cells were sonicated to detach the head from the tail, followed by sequential centrifugation to enrich each fraction (Supplementary Figure 4A). We analysed competitive mixing fractions at defined head-to-tail ratios (100/0, 80/20, 60/40, 40/60, 20/80, 0/100) across six donors, with five replicates per donor, and identified proteins following the expected abundance profile trends. The total number of protein groups identified was consistent across fractions, at approximately 8,700 each (Figure 3B). A Linear Models for Microarray (Limma) differential t-test-based analysis of relative protein abundances, visualised as a volcano plot, showed a major difference in protein intensities between the most divergent fractions (100/0 vs 0/100 head/tail) (Figure 3C), and the least difference between near-equal fractions (60/40 vs 40/60) (Supplementary Figure 4B). Known tail-localised proteins, including the CatSper complex, and head-localised acrosomal enzymes such as acrosin (ACR) were detected in each compartment as expected (Figure 3D), confirming the successful separation of the head and tail proteins. For each protein, we computed mean intensity across the five replicates of each head:tail mixing ratios (0:100 to 100:0) and used the Pearson correlation of this six-value profile against a monotonically increasing reference vector as a localisation score, ranging from-1 (tail behaviour) to +1 (head behaviour). Proteins scoring above +0.75 were classified as head-localised, below-0.75 as tail-localised, and the remainder as unassigned, yielding 2,629 head-and 2,624 tail-assigned proteins (Supplementary Figure 4D).

**Figure 3.**
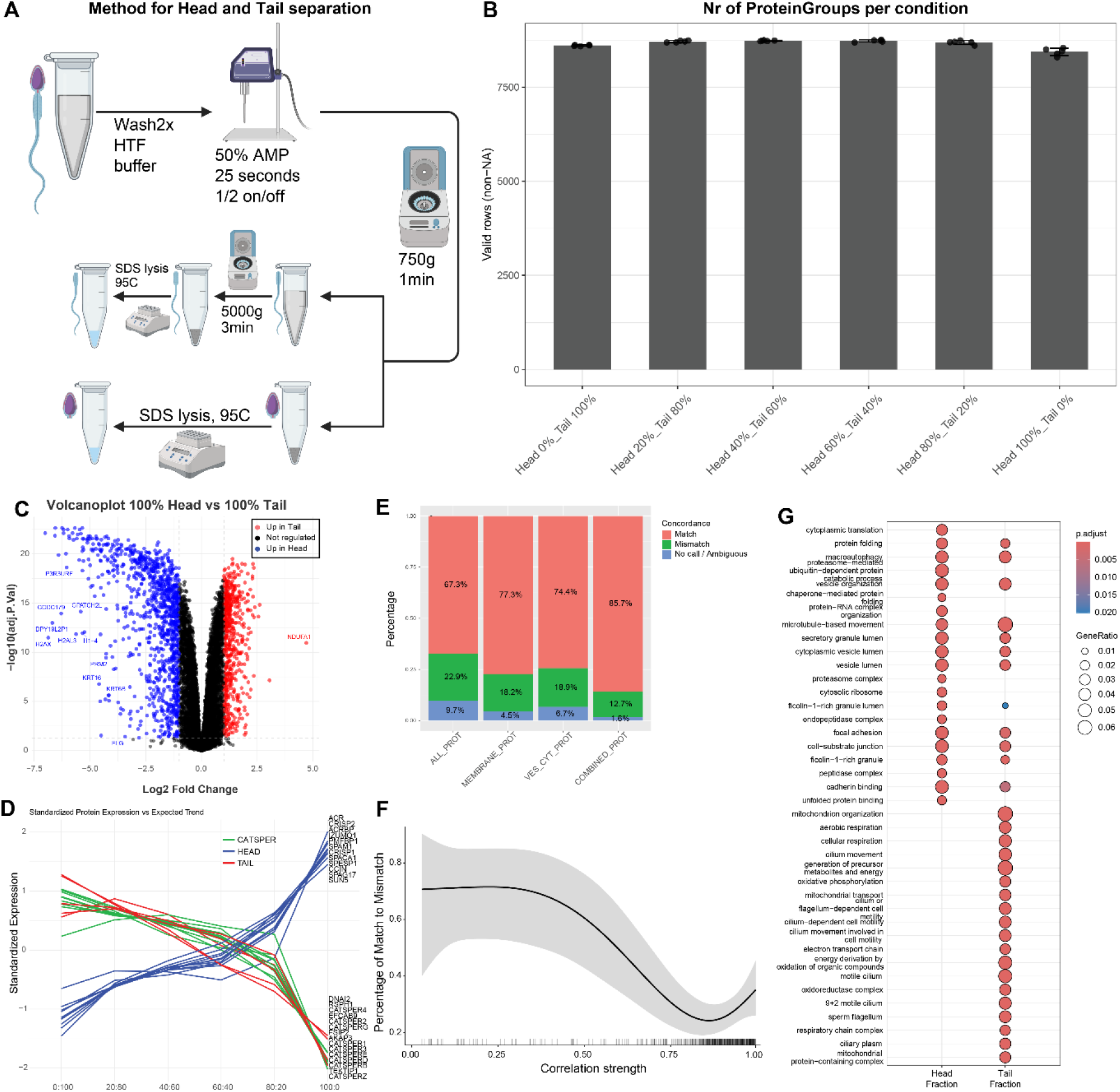
Spatial resolution of the sperm proteome into head and tail compartments. (A) Schematic of the head–tail separation workflow: washing, sonication to detach heads from tails, and sequential centrifugation to enrich each fraction prior to SDS lysis and digestion. (B) Total protein group identifications across competitive head-to-tail mixing fractions (100/0, 80/20, 60/40, 40/60, 20/80, 0/100); approximately 8,700 proteins per fraction. (C) Volcano plot of differential protein abundance between the 100/0 and 0/100 head/tail fractions. Proteins enriched in the head (red) and tail (blue) are highlighted. (D) Standardised intensity profiles of known head-and tail-localised proteins across the mixing fractions. CatSper complex components and other tail markers (green/red) decrease with increasing head content, while head markers increase. (E) Concordance between head/tail assignments from this study and Human Protein Atlas annotations. (F) Match rate (matches / [matches + mismatches]) as a function of absolute localisation score. (G) GO term enrichment of head-and tail-assigned proteins

To corroborate this, we integrated HPA compartment annotations (628 proteins), grouped into head (acrosome, equatorial segment, perinuclear theca, calyx) and tail (principal piece, end piece, annulus, connecting piece, flagellar centriole, midpiece, mitochondria) categories. We validated our 0.75 localisation-score threshold, chosen to balance stringency and coverage, against these annotations (Figure 3E). Overall concordance was 67.3%, with 22.9% mismatch and 9.7% no-call; restricting to membrane proteins improved this to 77.3%, excluding vesicular/cytoplasmic proteins to 74.4%, and the combined filter to 85.7% (Figure 3E). Lower localisation scores were associated with lower annotation confidence and higher mismatch rates (Figure 3F, Supplementary Figure 4F); although HPA annotations, being antibody-based, may themselves contain false localisations, we suspected this trend also reflected protein abundance, via signal saturation or loss at low abundance. A Wilcoxon test comparing matched and mismatched protein abundances confirmed a clear difference (p = 0.0119) (Supplementary Figure 4E). GO term analysis showed head-fraction proteins enriched for protein folding, vesicle organisation, macroautophagy, proteasome-mediated catabolism and cytoplasmic translation, and tail-fraction proteins enriched for cellular and aerobic respiration, mitochondrion organisation, cilium movement and microtubule-based movement, all well-established sperm tail processes (Figure 3G, Supplementary Figure 4C).

### Single-cell proteomics of individual sperm cells and multiple donors

We next explored the feasibility of performing single-cell sperm proteomics. Sperm cells are ∼50 µm long, but the head is only ∼4-5 µm, roughly 7-8× smaller than somatic cells, with an estimated protein content of only ∼2-5 pg, representing a challenge for low-input single-cell proteomics. Cells were isolated using an automated sample-preparation workflow (44) on a cellenONE robot, collecting 1-, 5-, 10-and 20-cell samples alongside blank controls. After filtering, the median number of protein groups differed substantially between donors (D1 and D3: ∼1,000; D2 and D4: ∼500 per single cell; Figure 4A). Identifications increased consistently with the number of single-sperm included while blanks remained virtually empty (Figure 4A), confirming genuine signal over background; 20-cell samples yielded ∼2,000 protein groups on average (Supplementary Figure 5A), with the same trend at the precursor level (Supplementary Figure 5B). Reproducibility was consistent across donors, with single-cell CVs of ∼40-50% decreasing to ∼13% at 20 cells (Figure 4B), and median log2 protein intensity was stable across most samples (∼10-17) (Supplementary Figure 5C).

**Figure 4.**
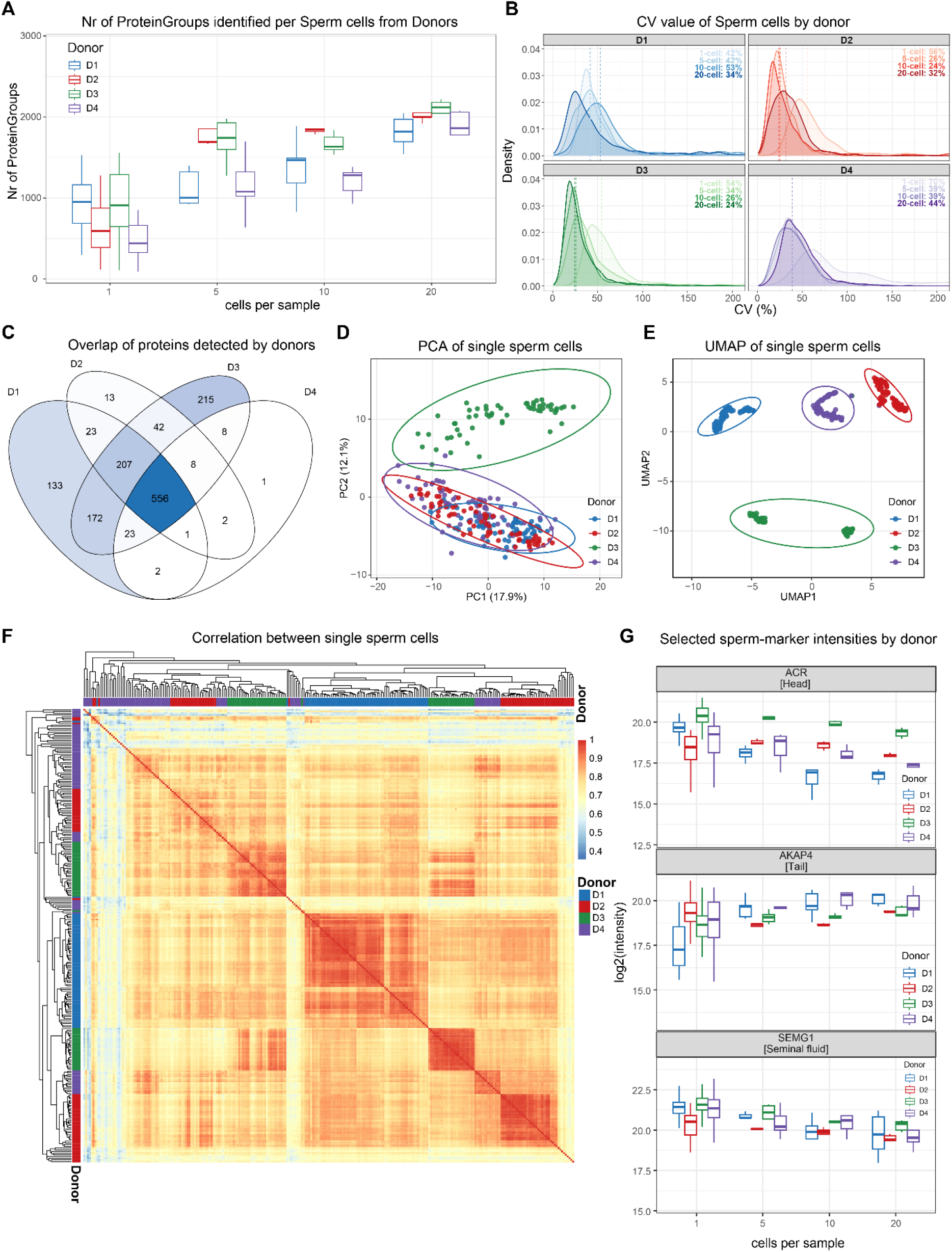
Single-cell proteomics of individual sperm cells from 4 donors. (A) Number of protein groups identified per single sperm cell and per 5-, 10-, and 20-cell inputs, separated by donor (D1–D4). (B) Density distributions of the coefficient of variation (CV, %) across single cells and 5-, 10-, and 20-cell inputs, shown per donor. Percentages indicate the median CV for each input condition. (C) Venn diagram showing the overlap of proteins detected across the four donors. (D) PCA of single sperm cells colored by donor, with 95% confidence ellipses per donor. (E) UMAP of single sperm cells colored by donor, with 95% confidence ellipses per donor. (F) Correlation heatmap between single sperm cells with hierarchical clustering; donor identity is annotated along the rows and columns. (G) log2 intensity of selected sperm-marker proteins (ACR [head], AKAP4 [tail], SEMG1 [seminal fluid]) across single cells and 5-, 10-, and 20-cell inputs, separated by donor.

To confirm that this donor structure did not depend on restricting the analysis to a shared protein subset, we computed a UMAP embedding from all 2,735 proteins detected in single cells, using pairwise-complete correlation distances so that no missing values were imputed. The same four donor clusters were recovered (Figure 4E). Correlation between single cells showed clear within-donor structure alongside clusters spanning all four donors (Figure 4F), with pairwise Pearson correlations between donors ranging from r = 0.849–0.900 (Supplementary Figure 5E). Tracking markers of the head (ACR), tail (AKAP4) and seminal fluid (SEMG1), we observed that ACR log2 intensity unexpectedly decreased with increasing cell number, AKAP4 intensity increased as expected, and SEMG1 remained stable (Figure 4G). The same opposing trends held across the top annotated proteins per compartment (Supplementary Figure 5F). We attribute the head-protein decrease to degradation during the prolonged sample handling required for cell sorting. The acrosome contains active serine proteases such as acrosin (45) and functional 26S proteasome subunits (46), and the processing of heads will likely trigger autocatalytic degradation of head-localised proteins. Tail structural proteins such as AKAP4, anchored within the disulfide cross-linked fibrous sheath (47) and outer dense fibres, resist proteolysis under the same conditions.

### Origin of the seminal fluid proteome

As proteins in the ejaculate originate from both spermatogenesis in the testes and from additions by the epididymis and accessory sex glands, we characterised their origin by analysing the seminal fluid proteome from vasectomised (n=13) and non-vasectomised (n=22) men, comparing pre-versus post-vas-deferens contributions. Close to 2,400 proteins were shared between groups (Figure 5A). The top 75 most abundant proteins in non-vasectomised men remained the most abundant after vasectomy (Supplementary Figure 6A, B), with the exception of FN1, CAMP, HSP90AA1, PTGDS and TMED5, which all decreased and are all expressed in the testis and epididymis (Human Protein Atlas, 48), the tissues cut off by the vasectomy. PCA showed clear separation between groups (Figure 5B); most proteins were shared with the pure sperm dataset, with 407 quantified only in seminal fluid (Supplementary Figure 6C). Limma analysis identified 294 significantly different proteins (205 higher in non-vasectomised, 89 higher in vasectomised men) (Figure 5C). GO over-representation showed reproductive-process enrichment in non-vasectomised men, consistent with loss of pre-vas deferens-derived proteins, and secretory-vesicle/vesicle-lumen enrichment in vasectomised men, reflecting a greater relative contribution of accessory-gland secretions (Figure 5D); STRING/MCL clustering further resolved functional groups among the vasectomised-specific (Supplementary Figure 6D) and non-vasectomised-specific (Supplementary Figure 6E) proteins. To deconvolute tissue origin, we assigned each protein a peak-expression tissue using HPA tissue-specificity scores, restricted to the four male genital glands (prostate, testis, epididymis, seminal vesicle), most were unassigned (2,279), with the remainder distributed across testis (74), epididymis (50), prostate (27) and seminal vesicle (9) (Supplementary Figure 6F). Weighting by log2 summed intensity rather than protein number, the seminal vesicle dominated the proteome despite contributing the fewest annotated proteins, consistent with it and the prostate providing the bulk of seminal fluid’s secretory volume (49). Its contribution, along with the epididymis, was lower in vasectomised men, while the prostate contribution was modestly higher (Figure 5E). The epididymal decrease is anatomically consistent, as the epididymis lies proximal to the vasectomy site. Testis-annotated proteins were essentially unchanged after vasectomy, with only three proteins that could be confidently assigned to a single tissue detected in non-vasectomised but not vasectomised men, and the seminal-vesicle share of total intensity was lower rather than higher. The unchanged testis and decreased seminal-vesicle contributions are harder to reconcile with anatomy, possibly reflecting inaccurate tissue annotation or the limited resolution imposed by the small number of gland-specific proteins and high donor-to-donor variability.

**Figure 5.**
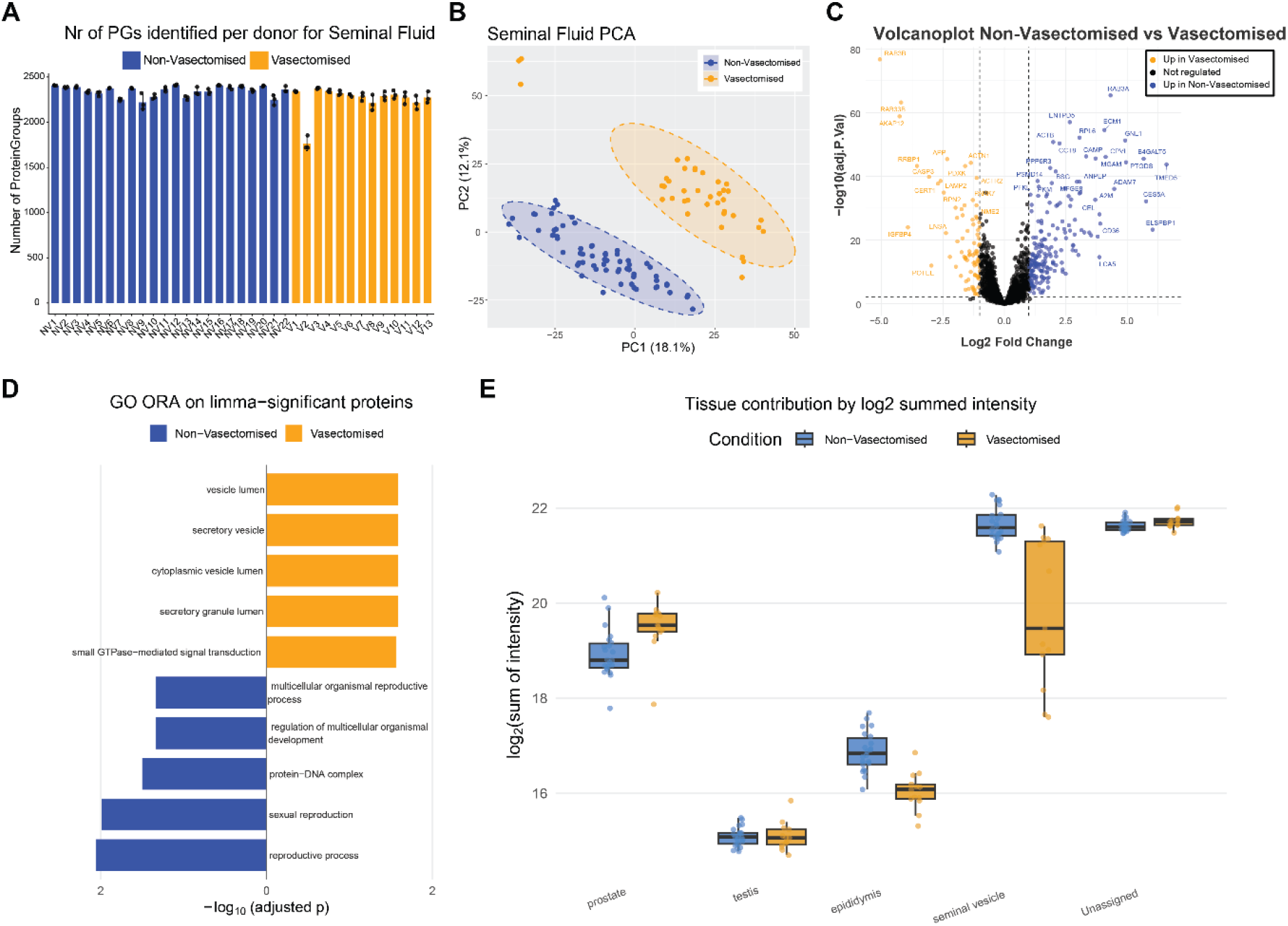
Proteomic characterisation of seminal fluid from vasectomised and non-vasectomised men. (A) Number of protein groups identified per sample for non-vasectomised (blue, n = 22) and vasectomised (orange, n = 13) men. (B) PCA of seminal fluid proteomes showing clear separation between non-vasectomised and vasectomised men. (C) Volcano plot of differential protein abundance between non-vasectomised and vasectomised groups. (D) GO term enrichment comparison of the most enriched terms in each group (non-vasectomised, blue; vasectomised, orange). (E) Tissue-of-origin contribution to the seminal fluid proteome by log2 summed intensity, split by condition.

### Proteomic signatures of total fertilisation failure

To translate our findings clinically, we investigated the molecular basis of Total Fertilisation Failure (TFF) (35) in couples with no zygote formation during in vitro fertilisation (IVF) attempts despite apparently normal semen and oocyte parameters. TFF affects 5-15% of IVF couples, but its causes remain poorly understood. While IVF renders some sperm functions less critical, zona pellucida penetration, sperm-oocyte fusion, energy metabolism and a degree of motility must remain intact. Because TFF can also originate from the oocyte, a sperm-centric interpretation must be made with caution. We analysed sperm from eight men with successful IVF fertilisation and five with TFF, all of whom later achieved pregnancy via intracytoplasmic sperm injection (ICSI). Given limited material from some patients, samples were analysed by low-input high-sensitivity LC-MS using Evosep Whisper flow gradients. PCA showed only partial separation with substantial overlap between groups (Figure 6A). Differential expression analysis nevertheless identified many proteins that differed between the groups (Figure 6B). We initially suspected sperm-egg fusion defects were responsible for the fertilisation failures, as this is a process susceptible to disruption in the context of TFF. To evaluate this, we examined the same proteins involved in gamete membrane fusion on spermatozoa (IZUMO1, SPESP1 and Syncytin-1) and O-glycosylation patterns (Tn and GALNT3) as done by Enoiu et al. (35), from which only patient ID31 showed the expected downregulation of IZUMO1, indicating dysregulated sperm-egg fusion (Figure 6C). Fusion-related dysregulation was therefore restricted to a single patient and does not account for the fertilisation failure in the other four, prompting us to look for changes shared across the cohort. After filtering for detection in at least half the samples of one group and reduced abundance in 4/5 TFF samples relative to IVF-success controls, we identified 54 proteins (Figure 6D, shown as z-scores relative to the eight IVF-success men). These were dominated by mitochondrial and metabolic proteins, including respiratory-chain subunits (MT-ND4, MT-ND5, COX6B1, ATP5MF) and mitochondrial carriers (SLC25A2, SFXN1, SLC30A1). Cytoscape analysis after MCL clustering and functional enrichment returned Mitochondrial Matrix as the top term (Figure 6E), suggesting that mitochondrial rather than fusion dysregulation likely underlies the fertilisation failure in these men.

**Figure 6.**
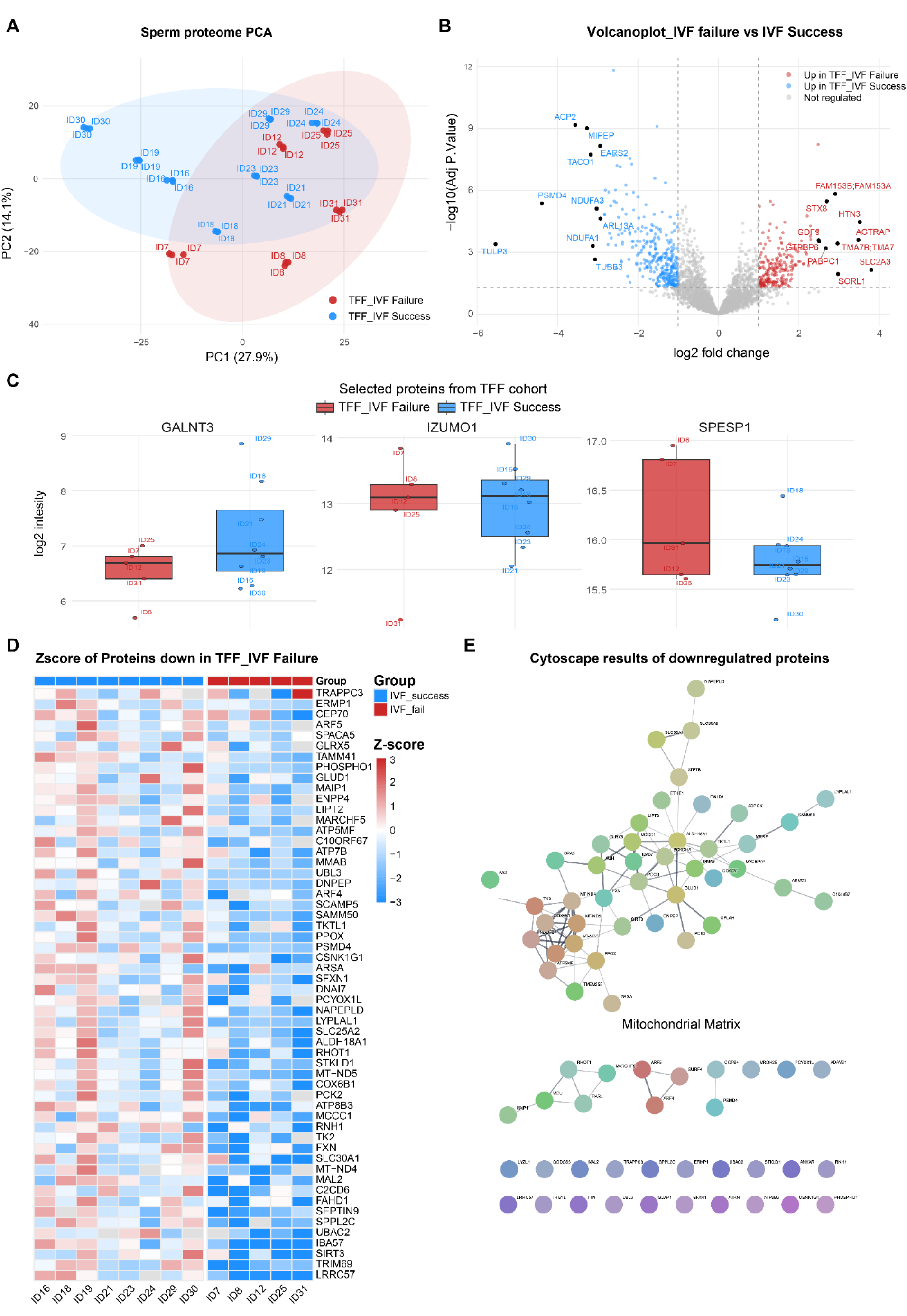
Sperm proteomic signatures of total fertilisation failure. (A) PCA of sperm proteomes from men whose partners underwent IVF, comparing IVF success (blue) and total fertilisation failure (TFF/IVF failure, red). (B) Volcano plot of differential protein abundance between the TFF/IVF failure and IVF success groups. (C) Log2 intensity of selected fertilisation-related proteins (GALNT3, IZUMO1, SPESP1) in the TFF/IVF failure (red) and IVF success (blue) groups. (D) Heatmap of the 54 proteins consistently reduced in the TFF/IVF failure samples relative to the eight IVF-success men, shown as z-scores and grouped by condition. (E) Cytoscape network of the downregulated proteins after MCL clustering and functional enrichment.

## Discussion

This study presents the most comprehensive proteome map of human spermatozoa to date, identifying 10,828 proteins through single-shot nDIA acquisition on an Astral mass analyser combined with off-line high-pH reversed-phase fractionation, a 16% increase over Kong et al. (15) and more than double of Greither et al. (21). The high correlation with Kong (r = 0.855) and the lower correlation with Greither (r = 0.589) are consistent with the different donors, acquisition strategies and software, while the conserved rank order of the most abundant proteins and recovery of 85% of the theoretical sperm proteome proposed by Pini et al. (14) support the identifications. Infertility affects approximately one in six adults worldwide (50), with male factors contributing to a substantial proportion of cases against a background of declining sperm counts (51), and the added depth obtained here provides biological insight and potential clinical applications in that context. We also observed differences in the sperm proteome between donation time points within the same men, suggesting that the proteome is not fixed and may respond to factors that vary between donations. The increasing sensitivity of mass spectrometers also makes it more likely that carryover or trace contamination will be detected. Residual contamination, to which the Astral is more sensitive (36), proteins introduced during sample preparation, or proteins adhering to the outer cell surface therefore cannot be fully excluded. This is relevant to the 2,655 proteins detected only in our dataset relative to Kong et al., which were enriched for processes characteristic of spermatogenic precursors rather than the mature cell. Some are likely remnants of spermatogenesis that are incompletely degraded or recycled during maturation (46, 52), and others may be delivered with organelle-derived membranes, as in the Golgi-derived proacrosomal vesicles that form the acrosome. Residual translational activity has also been reported in mature sperm (8, 53, 54), although this would account for only a subset of the terms. One way to potentially remove these proteins could be treatment with proteinase K to shave off extracellular proteins, as recently done for extracellular vesicles (55), though this would come at the cost of digesting some membrane proteins.

Repeated within-donor sampling of the sperm sncRNAome has shown that between-individual differences dominate over within-individual change across six monthly collections, with only a minority of transcripts behaving dynamically (56), although diet and other exposures can shift the profile within weeks (42, 43). The proteome showed a comparable architecture, with a conserved, largely mitochondrial core alongside a smaller fraction that changed between donations. This variable fraction may be the more informative one clinically. A proteome fixed at spermiation and invariant thereafter would offer little beyond a static description of a man, whereas one that changes between donations could report on current reproductive state and, in principle, on response to intervention. It also implies that a single ejaculate may not fully characterise an individual, and that repeated sampling should be considered in diagnostic use. Although we abstinence was held constant, we did not record diet or other exposures, nor standardise the interval from ejaculation to liquefaction and processing, so the causes of this variation remain to be established. Especially the liquefaction could potentially have an effect on the sperm proteome as this will lead to more degradation of sperm surface proteins. Therefore, in future studies this would need to be corrected for.

Our head-and-tail fractionation assigned subcellular localisation to a large fraction of the proteome (2,629 head, 2,624 tail proteins), with 85.7% concordance with Human Protein Atlas (37) annotations when restricted to membrane proteins and no vesicle proteins, consistent with soluble proteins redistributing during mechanical separation while membrane-associated proteins retain their native compartment. Localisation is especially informative in spermatozoa, which are transcriptionally and translationally silent and cannot re-localise or replace their proteins, so spatial position is close to a functional assignment: head proteins act in recognition, penetration and fusion, tail proteins in energy generation and motility. This is reinforced by the GO enrichment of newly localised proteins, quality control and autophagy in the head, mitochondrial and ciliary function in the tail. A compartment-resolved proteome can therefore link a molecular defect to a clinical phenotype class, mapping tail-restricted changes to motility disorders and head-restricted changes to acrosomal or fusion failure.

We further demonstrate the feasibility of single-cell sperm proteomics, recovering a median of ∼500–1,000 protein groups per cell (rising to ∼2,000 in 20-cell samples) with virtually empty blanks and CV decreasing as input increased, confirming genuine signal over background. Canonical markers showed an unexpected decrease in head-protein ACR intensity with increasing cell number, against rising tail AKAP4 and stable SEMG1, which we attribute to autocatalytic degradation during handling: the acrosome contains active serine proteases (45) and proteasome subunits (46), whereas tail structural proteins are anchored within the disulfide cross-linked fibrous sheath and outer dense fibres (47) and resist proteolysis. Because bulk measurements average over the very heterogeneity that characterizes a sperm cell population, an ejaculate being defined by its most competent cells rather than its mean, coupling single cell with spatial resolution would be powerful for sperm analysis. This is not yet sensitive enough for clinical use, but with continued advances in sample preparation and acquisition it should become tractable.

Comparing seminal fluid from vasectomised and non-vasectomised men, we found ∼2,400 shared proteins and 407 quantified only in seminal fluid relative to the pure sperm dataset; clear PCA separation and extensive differential abundance show how substantially pre-versus post-vas-deferens input reshapes the ejaculate. The marked decrease in FN1 after vasectomy is coherent, as it is secreted partly by the epididymis (57), whose contribution is lost. Tissue-of-origin deconvolution placed the seminal vesicle as the dominant contributor by intensity, consistent with its secretory volume (49), though few proteins could be confidently assigned to a single gland and high donor variability limited resolution. Two aspects of the deconvolution did not fit the annotated anatomy, as testis-annotated proteins were essentially unchanged after vasectomy and the seminal-vesicle contribution fell rather than rose, whereas removing pre-vas-deferens input should have produced the opposite in both cases. Part of this is likely technical, since HPA tau scores reflect enrichment rather than exclusivity, and only 9 proteins could be confidently assigned to the seminal vesicle and 74 to the testis, so the assignments are both approximate and sparse. The contributions are also relative and constrained to sum to one, so a lower seminal-vesicle share can follow from a rise elsewhere rather than from reduced secretion. Biologically, testis-derived proteins might also reach the ejaculate through the circulation rather than through the vas deferens alone, though this remains speculative, and the accessory glands may themselves be regulated by testicular and epididymal factors lost at vasectomy, so that their secretory output changes rather than a compartment being fully removed from, for example serum androgens are unaffected by the procedure (58). Continued passage of fluid past the occlusion cannot be excluded despite the confirmed absence of spermatozoa, which would allow testicular and epididymal proteins to keep reaching the ejaculate.

In the TFF cohort we initially expected sperm–egg fusion dysregulation, as proposed by the referring clinicians (35). The data did not support it: only patient ID31 showed the anticipated IZUMO1 downregulation. Instead, the 54 proteins consistently reduced across 4 of 5 TFF samples relative to IVF-success controls were dominated by mitochondrial and metabolic constituents, and MCL clustering returned Mitochondrial Matrix as the top term, pointing to mitochondrial rather than fusion dysfunction as a recurrent feature in this cohort. Because TFF can also originate on the oocyte side, this sperm-centric reading is provisional and would require parallel oocyte analysis to confirm. Repeated sampling of the same men would also help to determine whether this signature is stable or specific to the failed ejaculate.

Measuring the proteome across such different inputs helps us distinguish which proteins are truly sperm-intrinsic rather than co-isolated contaminants or potentially surface-bound. Single-cell data are the most stringent filter: with only an isolated spermatozoon in the well, the ∼500–1,000 proteins per cell (up to ∼2,000 pooled) are effectively free of seminal-plasma and accessory-gland input and therefore define a high-confidence core. A limitation here is that we will mostly detect the highest abundance proteins only due to the low input. Head and tail fractionation (2,629 and 2,624 proteins) adds spatial support, as proteins that reproducibly partition to a compartment are unlikely to be adventitious, and the vasectomised versus non-vasectomised comparison further distinguishes testis-and epididymis-derived from accessory-gland proteins. By contrast, the ∼10,000 proteins from bulk samples are the most permissive layer, also capturing carryover, preparation artefacts and surface-adhered seminal proteins, so the resource is best read as a confident sperm-intrinsic core surrounded by a lower-confidence, contaminant-prone periphery.

Taken together, this work provides the deepest human sperm proteome assembled to date, together with two approaches for further investigation mechanical head-tail fractionation, which assigns a compartment to thousands of previously unlocalised proteins, and single-cell acquisition, which establishes that individual spermatozoa are measurable. Mitochondrial proteins recur across these layers, forming the most conserved fraction between donors, partitioning to the tail, and dominating the proteins consistently reduced in total fertilisation failure. As a resource, this provides a starting point for male-infertility diagnostics; as a method, it points toward proteome measurement at the level of the individual cell, where the heterogeneity of an ejaculate is likely to reside.

## Supplementary

**Supplementary Figure 1.**
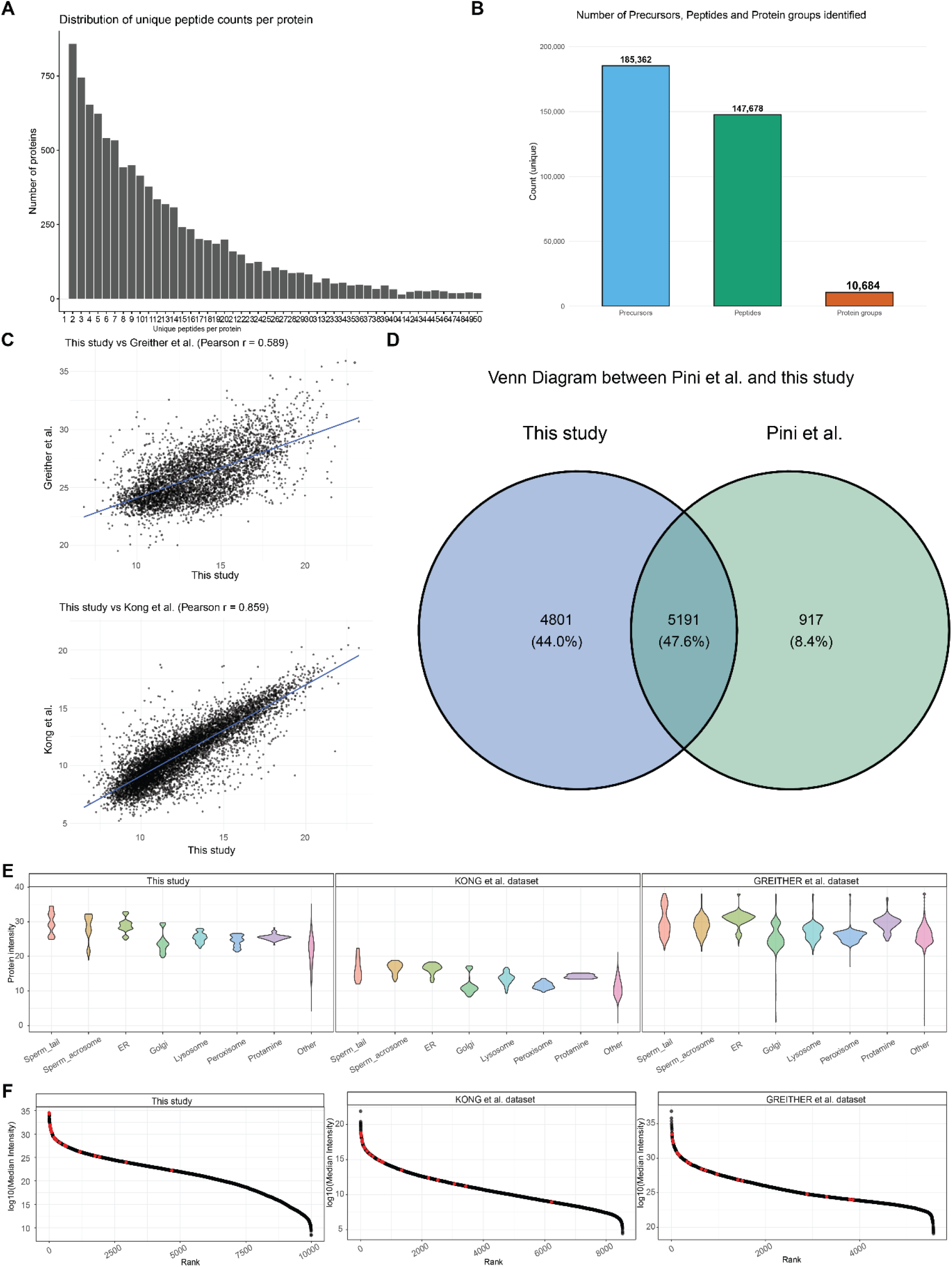
Extended validation of the human sperm proteome depth. (A) Distribution of unique peptide counts per protein across the dataset, reflecting proteome coverage. (B) Total numbers of precursors, peptides and protein groups identified. (C) Pearson correlation of protein intensities between this study and the previously published Kong et al. (r = 0.859) and Greither et al. (r = 0.589) datasets. (D) Venn diagram comparing the current dataset against the sperm proteome defined by Pini et al., after collapsing redundant and unannotated entries to a non-redundant gene set via UniProt. (E) Abundance distribution of proteins annotated to organelles not expected in mature sperm cells (e.g. ER and Golgi apparatus), examined as a contamination quality check. (F) Abundance rank of the most highly expressed HeLa cell proteins within the current sperm dataset, compared against previously published sperm proteome datasets.

**Supplementary Figure 2.**
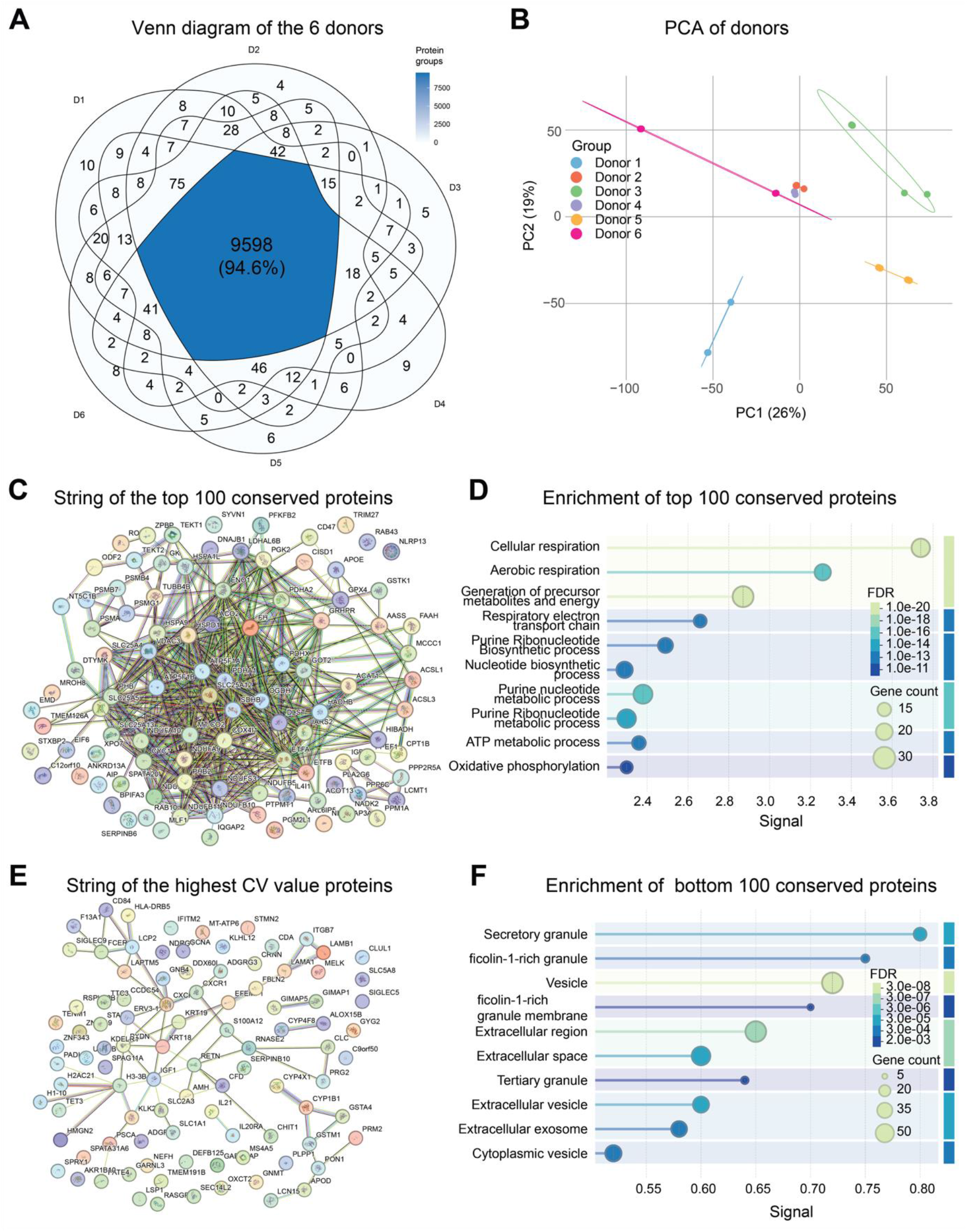
Extended analysis of inter-donor variability and temporal stability. (A) Venn diagram of the overlapping proteins from the 6 donors. (B) Principal component analysis (PCA) of the sperm proteome across the six donors (D1–D6), with samples coloured by donor. (C) STRING protein-protein interaction network of the 100 proteins with the lowest inter-donor CV. (D) GO term enrichment of the 100 lowest-CV proteins using the STRING function (E) STRING network of the 100 proteins with the highest inter-donor CV. (F) GO term enrichment of the 100 highest-CV proteins, dominated using the STRING function

**Supplementary Figure 3.**
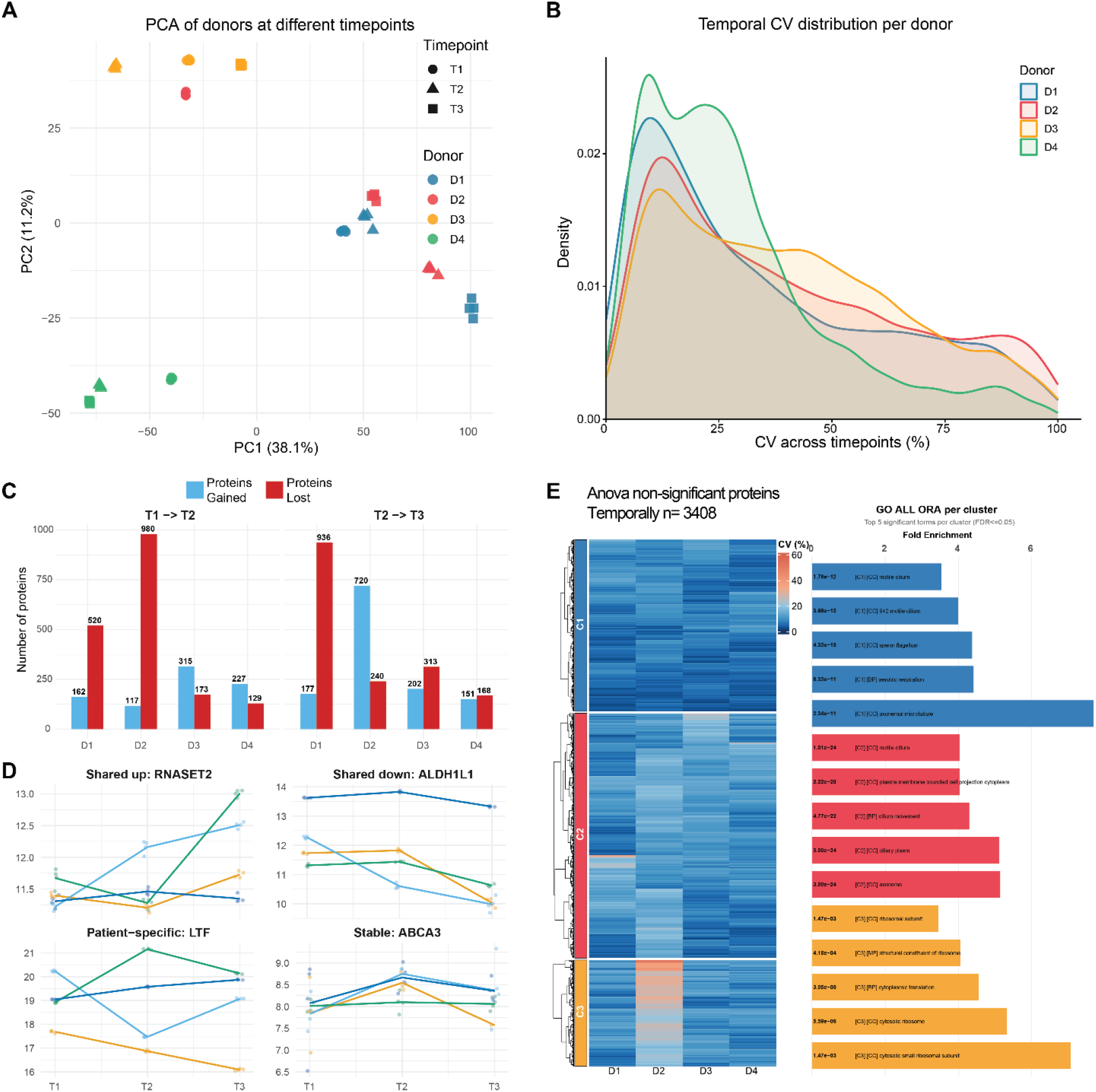
Temporal stability of the sperm proteome across donation time points. (A) PCA of all donor samples collected at three donation time points (T1, T2, T3), coloured by donor (D1–D4) and shaped by time point. (B) Density distribution of the per-protein coefficient of variation (CV) calculated across time points for each donor. (C) Number of proteins gained (blue) and lost (red) between consecutive time points (T1→T2 and T2→T3) for each donor, quantifying protein turnover over time. (D) Log2 intensity trajectories across T1, T2 and T3 for representative proteins from four temporal categories: shared increase (RNASET2), shared decrease (ALDH1L1), patient-specific (LTF) and stable (ABCA3), with lines coloured by donor. (E) Heatmap of the 3,408 proteins not significantly different across time points by ANOVA, coloured by per-donor CV and grouped by hierarchical clustering into three clusters (C1–C3) across donors D1–D4, with GO over-representation analysis of each cluster (top five terms, FDR ≤ 0.05) shown on the right.

**Supplementary Figure 4.**
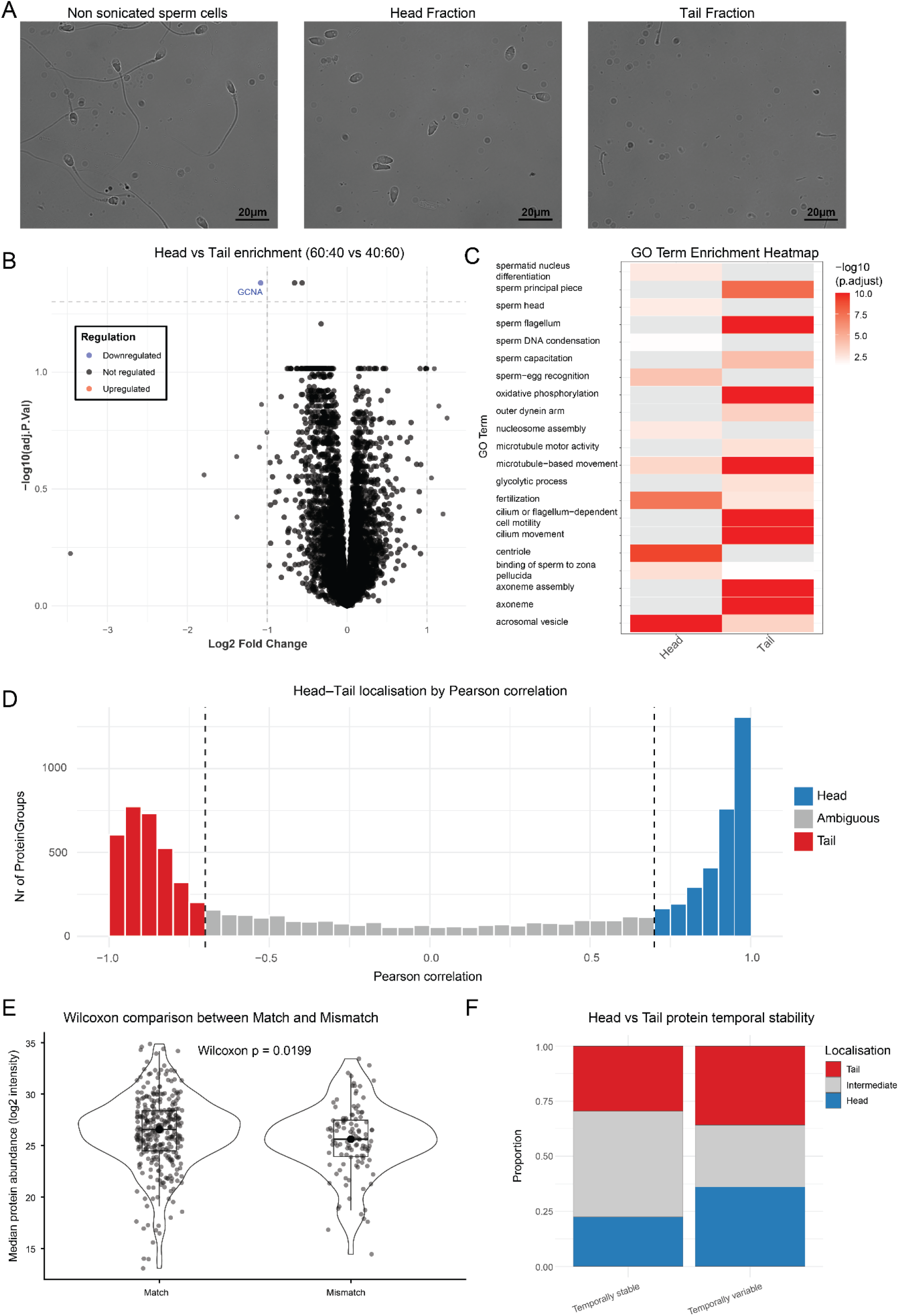
Extended validation of sperm head and tail compartment assignments. (A) Images of sperm cells, on the left non sonicated sperm cells, in the middle the head fraction after sonication, and on the right the tail fraction. (B) Volcano plot of differential protein abundance between the 60/40 and 40/60 head/tail competitive mixing fractions, showing minimal differences between near-equal fractions. (C) GO term enrichment for selected GO terms comparing the Head and Tail fraction specific groups. (D) Classification of proteins as head, tail, or ambiguous, based on the Pearson correlation of their abundance profile to an idealized head or tail gradient. (E) Violin plot with Wilcoxon test (p = 0.0119) comparing protein abundances of matched versus mismatched proteins relative to HPA annotations, showing that mismatched proteins tend to be of lower abundance. (F) Proportion of head, intermediate and tail-localised proteins among the temporally stable and temporally variable proteins.

**Supplementary Figure 5.**
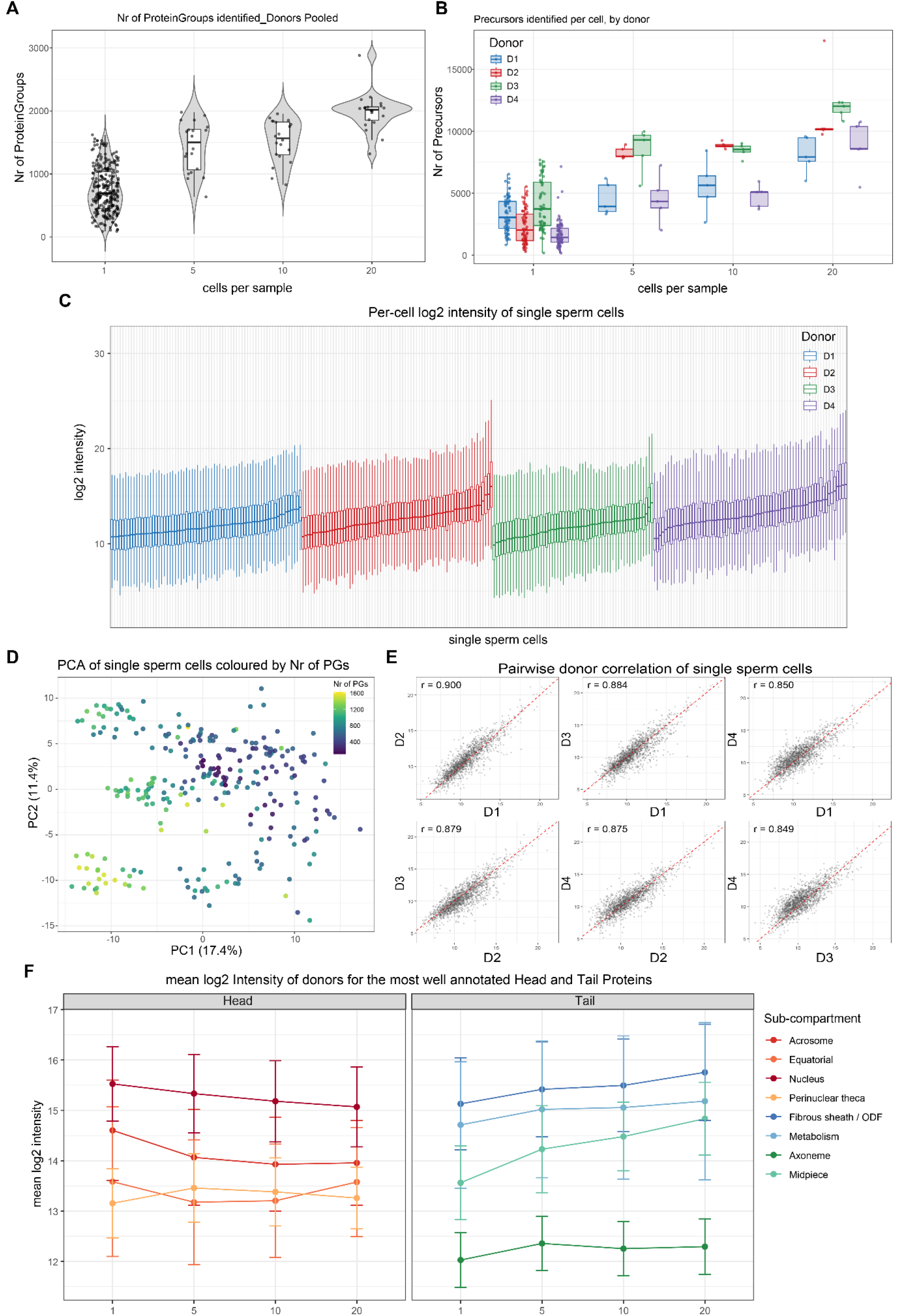
Extended single-cell sperm proteomics data. (A) Number of PGs identified per sample across inputs of 1, 5, 10 and 20 cells, pooled across donors. (B) Number of precursors identified per sample across the same inputs, shown separately for each donor. (C) Per-cell log2 intensity distributions for individual single sperm cells, coloured by donor. (D) PCA of single sperm cells coloured by the number of protein groups identified per cell. (E) Pairwise Pearson correlations of protein intensity between donors for single sperm cells (all pairwise combinations of D1–D4). (F) Mean log2 intensity across inputs (1, 5, 10 and 20 cells) for the best-annotated head and tail proteins, grouped by sub-compartment.

**Supplementary Figure 6.**
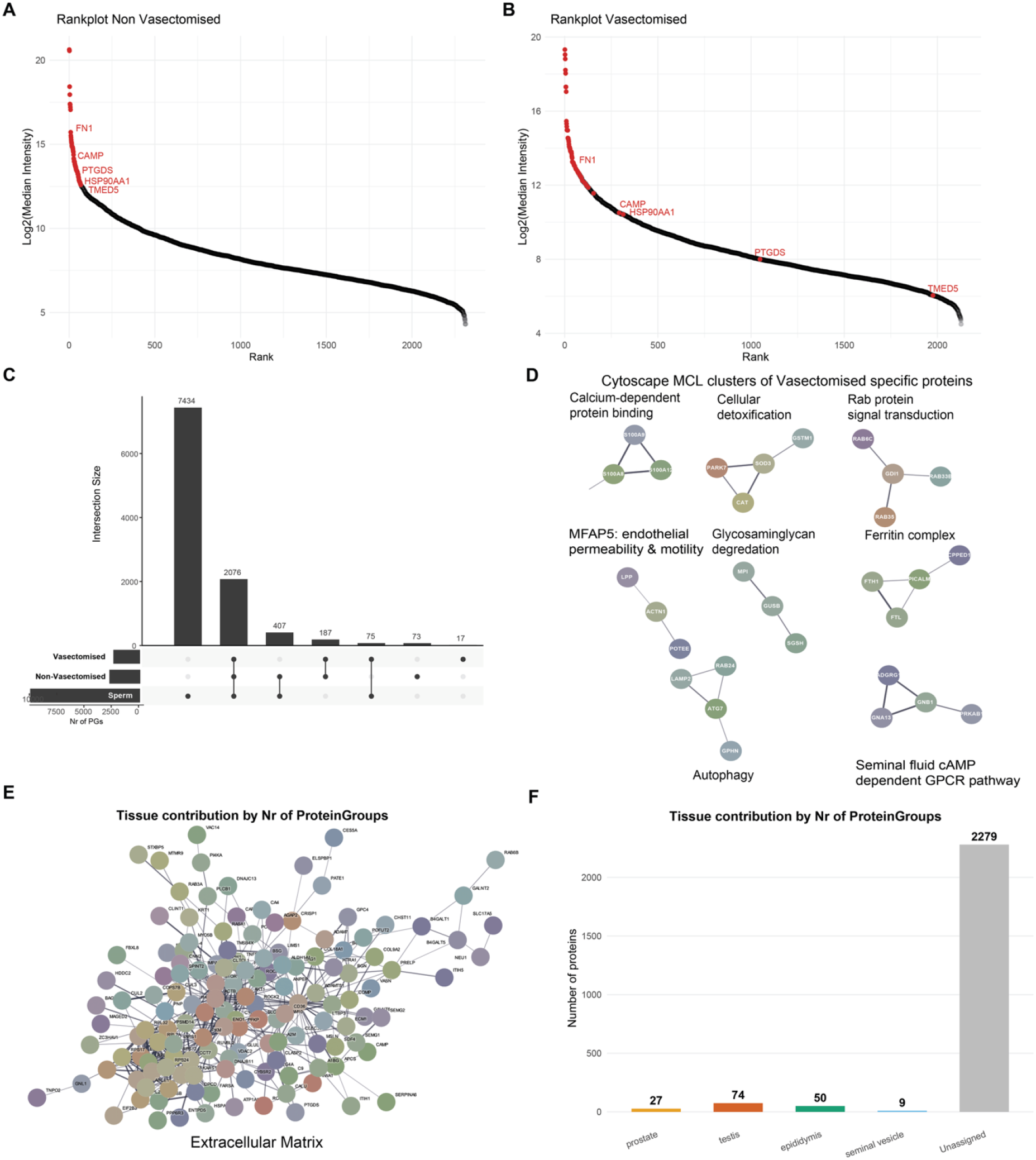
Extended characterisation of the seminal fluid proteome. (A) Rank plot of the vasectomised group, highlighting the top 75 most abundant proteins, and further labelling the ones that change markedly in rank between the Vasectomised and Non-Vasectomised group. (B) Rank plot of the non-vasectomised group, highlighting the top 10 most abundant proteins. (C) Upset plot comparing the ProteinGroups from the sperm protein, seminal fluid of vasectomised men, and seminal fluid of non-vasectomised men. (D) Cytoscape analysis of the ProteinGroups unique to the Vasectomised samples, followed by MCL clustering. (E) Cytoscape analysis of the ProteinGroups unique to the non-Vasectomised samples. (F) Barplot showing the Nr of proteins from the Seminal fluid that are assigned to specific tissues from the HPA annotation.

## Author Contributions

L.H. prepared samples, performed proteomics experiments, analysed the resulting data, and wrote the first version of the manuscript. L.S. performed the single sperm cell analysis. A.A.R., M.R.P., S.Z. and K.A. provided all sperm samples and critically evaluated the results. T.S.B. and J.V.O. critically evaluated the results. All authors read, edited, and approved the final version of the manuscript.

## Acknowledgements

Work at The Novo Nordisk Foundation Center for Protein Research (CPR) is funded in part by donations from the Novo Nordisk Foundation (NNF14CC0001, NNF24SA0098829, and NNF21OC0072070). This project was supported by a center-of-excellence grant from the Danish National Research Foundation to Copenhagen Center for Glycocalyx Research (DNRF196). This project was also supported by a generous grant from the Danish Agency of Higher Education and Science to establish the PLATO research infrastructure: Danish National Mass Spectrometry Platform for Proteomics and Biomolecular Imaging (grant no. 5229-00012B) and the Independent Research Fund Denmark Medical Sciences Instrument DFF-Research Project 1 (2034-00445 A).

## Ethical Approvals

All the samples used in this study are covered by the following ethical approvals H-19089581 (healthy donor cohort), and F-23073786 (seminal fluid cohort), H-17012149 (TFF cohort).

## Data Availability

Raw mass spectrometry data generated in this study have been deposited to the ProteomeXchangeConsortium via the MassIVE partner repository with the dataset identifiers PXD083465.

## Declaration of generative AI and AI-assisted technologies in the writing process

During the preparation of this work, the authors used Perplexity and Claude in order to improve language and readability. Additionally, it was used to generating code to be used in RStudio. After using these tools, the authors reviewed and edited the content as needed and take full responsibility for the content of the publication.

